# Biology-inspired data-driven quality control for scientific discovery in single-cell transcriptomics

**DOI:** 10.1101/2021.10.27.466176

**Authors:** Ayshwarya Subramanian, Mikhail Alperovich, Yiming Yang, Bo Li

**Author notes:** Equal contribution (Ayshwarya Subramanian, Mikhail Alperovich).

## Abstract

Quality control (QC) of cells, a critical step in single-cell RNA sequencing data analysis, has largely relied on arbitrarily fixed data-agnostic thresholds on QC metrics such as gene complexity and fraction of reads mapping to mitochondrial genes. The few existing data-driven approaches perform QC at the level of samples or studies without accounting for biological variation in the commonly used QC criteria. We demonstrate that the QC metrics vary both at the tissue and cell state level across technologies, study conditions, and species. We propose data-driven QC (*ddqc*), an unsupervised adaptive quality control framework that performs flexible and data-driven quality control at the level of cell states while retaining critical biological insights and improved power for downstream analysis. On applying *ddqc* to 6,228,212 cells and 835 mouse and human samples, we retain a median of 39.7% more cells when compared to conventional data-agnostic QC filters. With *ddqc*, we recover biologically meaningful trends in gene complexity and ribosomal expression among cell-types enabling exploration of cell states with minimal transcriptional diversity or maximum ribosomal protein expression. Moreover, *ddqc* allows us to retain cell-types often lost by conventional QC such as metabolically active parenchymal cells, and specialized cells such as neutrophils or gastric chief cells. Taken together, our work proposes a revised paradigm to quality filtering best practices - iterative QC, providing a data-driven quality control framework compatible with observed biological diversity.

## Introduction

Single-cell RNA sequencing (scRNA-seq) offers unprecedented resolution into cell biology by characterizing the individual cells within a biological sample of interest. Quality control (QC) of the cells is a critical first step in any scRNA-seq data analysis, which typically takes place after alignment of the sequencing reads to the reference genome (or transcriptome), and generation of the cell-by-gene matrix of gene expression counts. The goal of such QC is to remove “poor quality” cells, based on QC metrics such as the number of genes detected or (“gene complexity” or “transcriptional diversity”), the number of unique molecular identifiers (UMIs) recovered (typical for droplet based methods), and the fraction of mitochondrial and ribosomal protein genes [1]. The guiding motivation is that tissue dissociation techniques stress the cells and as cells die, transcription tapers off, cytoplasmic transcripts are degraded, and mitochondrial transcripts dominate [2]. Thus, low complexity of genes and high mitochondrial read content have been used as a proxy for identifying poor quality cells (or droplets with ambient RNA). As a corollary, high gene complexity has been used as a proxy for doublets or multiplets in droplet based sequencing [3]. While specialized computational strategies have been developed for specific tasks such as doublet identification [4–6], ambient RNA correction [7–9] or empty droplet removal [10], the standard practice in cell QC is to filter out cells by setting arbitrarily defined thresholds on the QC metrics. Widely used pipelines [11,12] by default, set a flat filter on the QC criteria for each sample or sets of samples analyzed, agnostic of the dataset and biology under study.

Although widely used, data-agnostic QC filters do not account for the fact that variation in the commonly used QC metrics may also be driven by biology (in addition to technical factors). For example, mitochondrial transcript abundance is dependent on cellular physiology [13], and metabolically active tissues (e.g. muscle, kidney) have higher mitochondrial transcript content [14,15]. Ribosomal protein gene expression has also been shown to vary by tissue [16] in human adults and mice [17]. Although biological variability in ribosomal protein gene expression has been reported [18], ribosomal protein gene expression is often conflated with technical artifacts or housekeeping transcription activity during analysis. Within each tissue, compartments and cell types may show further variability in these QC attributes. For example, the total number of genes expressed (gene complexity) varies with both cell type and cell state as seen during stages of mouse and human development [19]. Expression profiles also vary with progression through the cell cycle [20] or changes in cell volume [21]. Further, specific biological conditions or perturbations can lead to differences in these QC measures. For example, naive poised T-cells are known to have higher ribosomal content [22,23], as are malignant cells [24]. Activated lymphocytes such as Innate Lymphoid Cells (ILCs) [25] have greater transcriptional diversity, in an activation and condition dependent manner. Thus, the commonly used QC metrics can exhibit widespread biological variability bringing to the center the biological context of the study.

The importance of calibrating QC for the mitochondrial read fraction based on the mouse or human tissue of origin has been highlighted [26], however the proposed upper limit of 5 or 10% was largely based on existing data at the time of the study. Newer technologies (e.g. 10x v3 chemistry) may need a variable cutoff for mitochondrial read fraction [27]. The *scater* package [28] encourages the use of diagnostic plots and sample specific QC. More recently, probabilistic mixture modeling has been favored for data-driven quality control at the level of samples or sample sets, either in combination with other QC approaches [15] or standalone as in miQC [29]. However, no approach performs quality control explicitly taking into account the biological variability of QC metrics at the cell type or cell state level.

Here, we survey the variability of QC metrics across diverse scRNA-seq datasets at the tissue and cell state level, demonstrate the need for a data-driven quality control approach that accounts for the biological variability of QC metrics at the level of cell states, and present a framework for data-driven QC (*ddqc)*, inspired by unsupervised approaches in single-cell analysis, that performs adaptive quality control while retaining biological insights. Finally, we demonstrate that *ddqc* retains cell types that are lost by conventional QC, expanding existing cellular taxonomies for tissues, and offering an opportunity for further exploration and biological discovery.

## Results

### Survey of QC practices suggests a need for data-driven QC

To study existing QC practices in cell filtering, we sampled 107 research papers (**Methods**) with publication dates between 2017 and 2020, and focusing on analysis of scRNA-seq data generated across a range of technologies (3’ 10x V2 and 3’ 10x V3, Smartseq2, Drop-seq, mCEL-Seq2, Dronc-seq, MIRALCS, Microwell-seq) and in two species (mouse and human), and summarized the QC practices adopted (**Table S1).** The most commonly-used QC metrics were the number of genes detected, the number of UMIs counted, and the fraction of reads mapping to mitochondrial or ribosomal protein genes. A greater number of the studies (**Table 1**) that applied cell QC on specific metrics used data-agnostic QC filters, usually set at 5-10% for fraction of mitochondrial reads (86% or 73 papers), and 500 for gene complexity (86.5% or 77 papers). QC filter thresholds also varied with protocols (cells or nuclei) or technology (10x v3 vs v2 chemistry)[27]. Some studies excluded ribosomal protein or mitochondrial genes altogether or had a cutoff on the fraction of ribosomal genes [30]. Custom QC metrics were also adopted such as transcriptome mappability rates to exon vs non gene bodies [31] or the fraction of reads mapping to housekeeping [32] or select other genes such as *KCNQ1OT1[33]*, actin [34] or Hemoglobin [35]. Some studies had incorporated custom data-driven approaches including probabilistic mixture models [15,36] or sample [36,37] and study specific filters [38], suggesting the awareness and need for generalizable data-driven QC approaches. However, the majority of papers continue to use data-agnostic filters.

**Table 1.** Summary of QC survey. Quality control (QC) is typically performed for the following metrics: nUMI, nGenes, %mito (sometimes % ribo), and the minimum number of cells in which a gene is present. Additionally, empty droplets or multiplets may be detected and removed, and ambient RNA accounted for. We categorized studies into 4 groups: (1) Data-agnostic fixed threshold: The majority of the studies use a single threshold - for example <10% mitochondrial transcripts. (2) Many fixed thresholds: Sometimes, different fixed cutoffs are used for different samples within one study. (3) Sometimes studies regress out or remove mitochondrial and ribosomal genes. (4) Study-level threshold: There are also “data-driven” cutoffs (Data-Driven study-level threshold) - for example within 2 SDs from the median performed <u>on a per-sample</u> <u>basis</u>. Also, custom cut-offs are cutoffs that are very specific to the research in which they are used.

| Metric\QC type | Some QC | Data-agnostic fixed threshold (% of filtered) | Many fixed threshold | Removed | Data-Driven study-level threshold | Custom | No Filtering |
| --- | --- | --- | --- | --- | --- | --- | --- |
| nCounts | 65 | 48 (73.8%) | 5 (7.7%) | 0 (0%) | 11 (16.9%) | 1 (1.5%) | 42 |
| nGenes | 89 | 72 (80.8%) | 5 (5.6%) | 0 (0%) | 12 (13.5%) | 0 (0%) | 18 |
| nCells | 41 | 35 (85.4%) | 3 (7.3%) | 0 (0%) | 2 (4.9%) | 1 (2.4%) | 65 |
| %Mito | 85 | 69 (81.2%) | 5 (5.9%) | 4 (4.7%) | 6 (7.1%) | 1 (1.2%) | 22 |
| %Ribo | 7 | 2 (28.6%) | 0 (0%) | 3 (42.8%) | 2 (28.6%) | 0 (0%) | 100 |

| Empty Droplets | Doublets/Multiplets | Ambient RNA |
| --- | --- | --- |
| 4 | 17 | 6 |

### Across species and technologies, QC metrics vary by tissue

To systematically investigate if scRNA-seq data generated by commonly used technologies retains tissue and cell type specificity of the QC metrics, we profiled QC statistics by tissue and cell type on large public datasets after minimal basic QC (**Methods**). We surveyed 5,261,652 cells from 498 samples and 47 human tissues across 34 studies, and 966,560 cells from 337 samples and 37 mouse tissues across 5 studies (**Methods, Table S2**). We examined 8 human tumor types across protocols (fresh cells/scRNA-seq vs frozen nuclei/snRNA-seq) and droplet chemistries (10x v2 vs 10x v3) [27]. A subset of the studies (*Tabula muris* 10X, *Tabula muris* Smartseq2; Microwellseq mouse and human; *Tabula senis*) had both uniformly generated and processed datasets, while others (PanglaoDB) were generated in independent studies but uniformly processed. The mouse *Tabula muris* dataset was particularly convenient having data generated from both 3’-end droplet based sequencing (10X, (**Fig S1A,C,E**)) and full-length RNA plate based Smartseq2 techniques (**Fig S1B,D,F**) from the same samples, and processed uniformly using the same reference and computational pipelines.

We found a tissue specific (**Fig 1**) trend for the QC metrics across studies. In general, we found variation by tissue for proportion of mitochondrial reads (**Fig 1A,B**) within the same study regardless of the technology used (*Tabula muris* 10X, *Tabula muris* Smartseq2; Microwellseq mouse and human) with some tissues emerging as having higher mitochondrial content (e.g. kidney, colon, heart, liver etc). The tissue-specific ordering of mitochondrial reads seen in [13] was most faithfully recapitulated by the Smartseq2 dataset (**Fig S1B**) with kidney, colon, cerebellum and heart having the highest mitochondrial load. Differences in the gene complexity (**Fig 1C,D**) and the percent of ribosomal protein genes (**Fig 1E,F**) were also observed among tissues. Across both technologies, the tongue had the highest mean gene complexity (**Fig S1C,D**), with the mean percentage of ribosomal protein reads being higher in the 10X dataset (**Fig 1E**). Trends were generally also maintained with age (Tabula senis 30m, **Fig S2A,C,E**). When compared to frozen tumor nuclei, the gene complexity was higher for cells (**Fig S2D**). Further, within each tissue, multiple density modes were evident (**Fig 1**) for the QC metric studied. Finally, we note that the summary statistics of the QC metrics can vary by the experimental condition (technology and study) even for the identical tissue.

**Figure 1:**
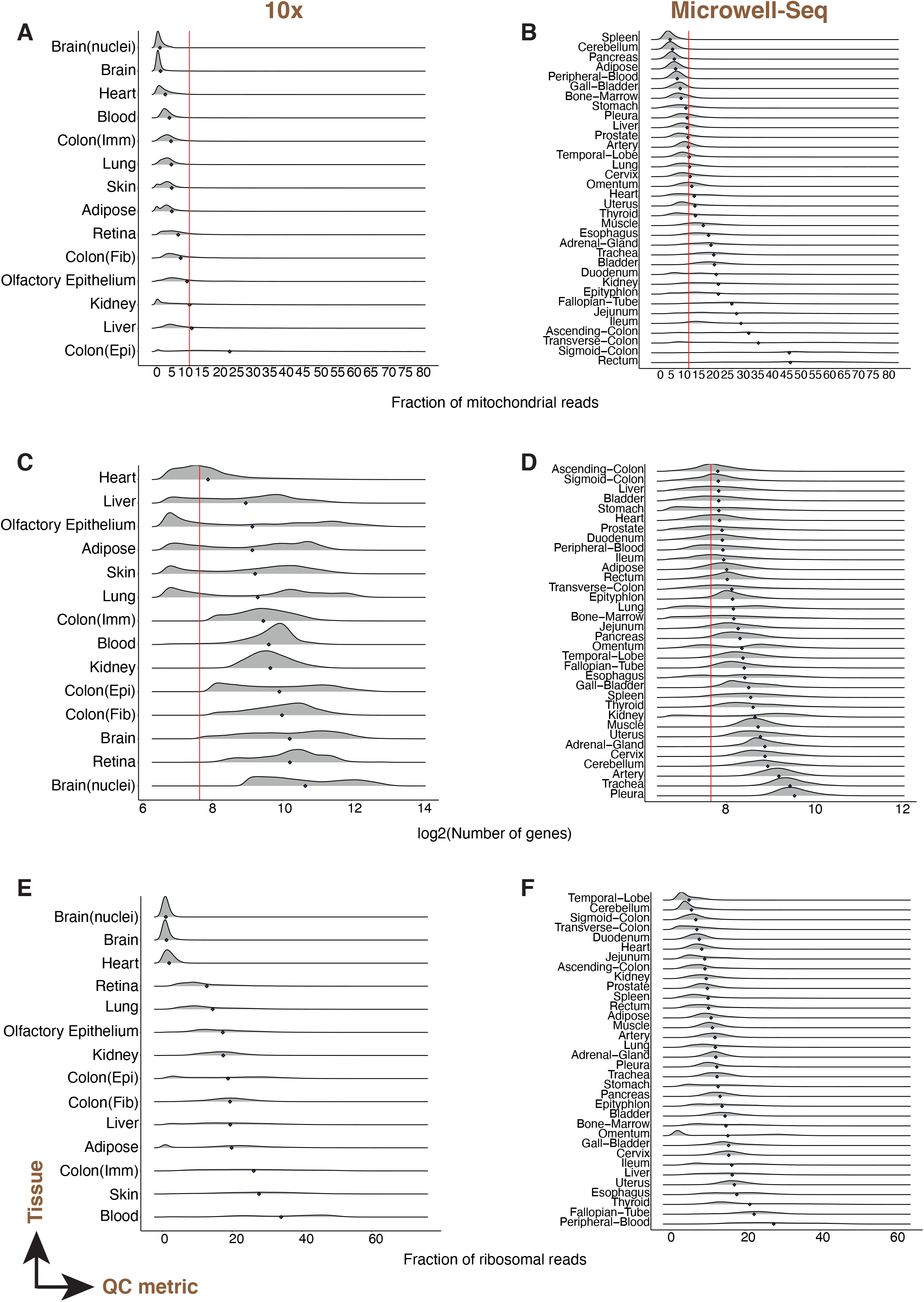
QC metrics vary by tissue: Fraction of mitochondrial reads (A,B), gene complexity (C,D) and percentage of ribosomal protein genes (E,F) per cell across human tissues and technologies. Various human tissue scRNAseq datasets generated by 10X droplet-based (A,C,E) and Microwellseq(B,D,F) technologies. Each row in a panel is a density curve with the mean represented by a blue diamond. Red lines indicate conventional threshold values set at 10% for percentage of mitochondrial reads, and 200 for gene complexity.

### Across species and technologies, QC metrics vary by cell-type within a tissue

We next assessed cell state or cell subset-specific QC attribute differences within tissues by uniformly processing all datasets (starting with the gene expression count matrices) to derive clusters within each tissue without applying standard QC cutoffs (**Methods**). However, many publicly available datasets did not come with assigned cell-type annotations. To uniformly predict biological annotations to the cell clusters, we devised a heuristic score function leveraging the top differentially expressed genes in a cluster, and the PanglaoDB [39] database of marker genes to assign the most probable cell type annotation. We tested the annotation strategy on 4 mouse (*Tabula muris* Smartseq2, *Tabula muris* 10X, *Tabula senis* 24 months, *Tabula senis* 30 months) and 1 human (Human Tissue Atlas) datasets which had partial annotations provided by the authors. On these data, our heuristic approach had an accuracy of 80.2% and 92.1% for cluster annotations in human and mouse data respectively (**Table S3**, **Methods**). We applied our heuristic approach to all test datasets and then examined trends of the QC metrics among cell states within tissues. As case studies, we manually verified annotations, and describe examples for murine (**Fig S2**) and human tissues (**Fig 2**).

**Figure 2:**
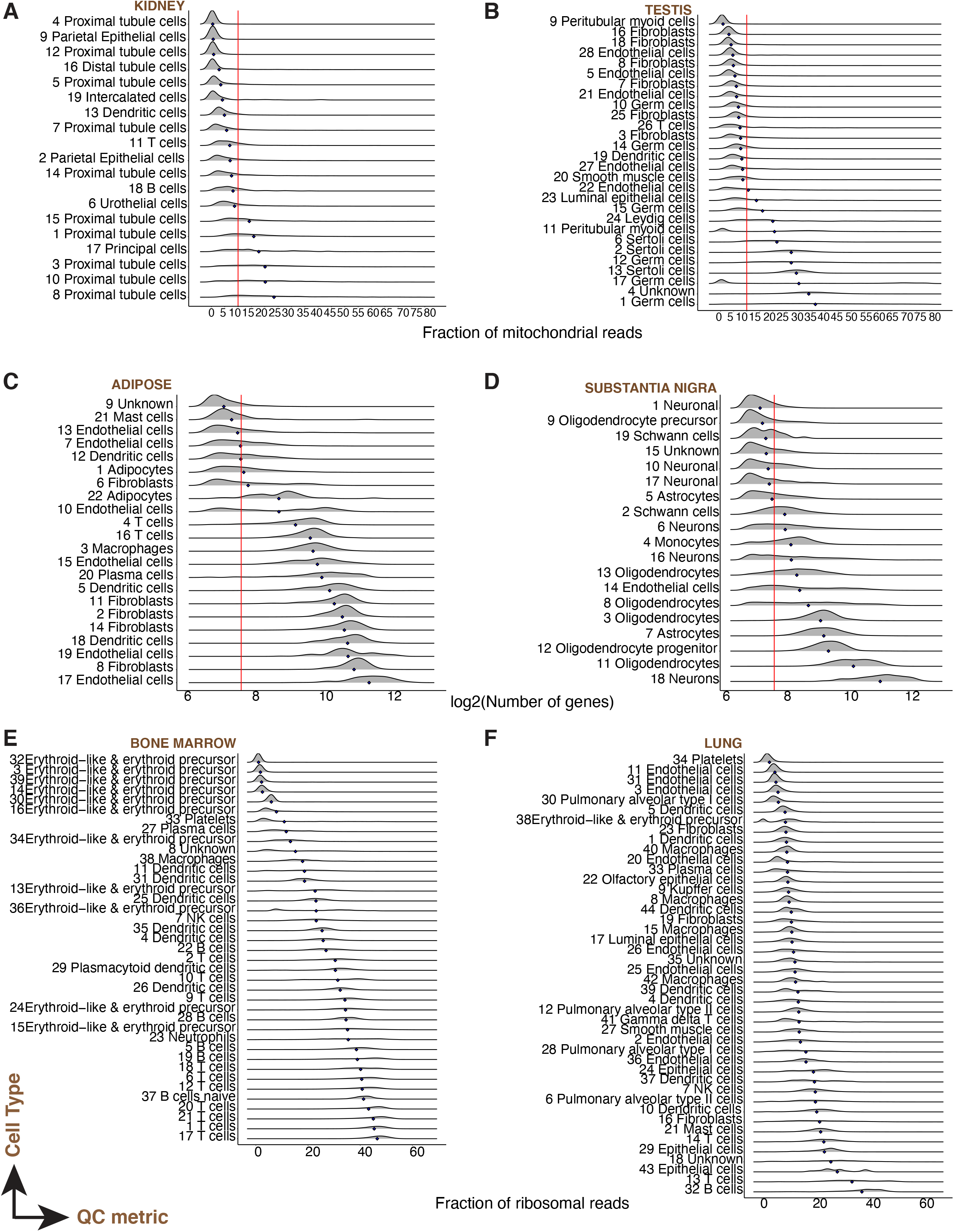
QC metrics vary by cell-type: Fraction of mitochondrial reads (A,B), gene complexity (C,D) and percentage of ribosomal protein genes (E,F) per cell across cell types of various human tissues: kidney (A), testis (B), adipose (C), Substantia Nigra (D), Bone Marrow (E) and Lung (F). All scRNA-seq data was generated using the 10X droplet-based technology. Each row in a panel is a density curve with the mean represented by a blue diamond. Red lines indicate conventional threshold values set at 10% for percentage of mitochondrial reads, and 200 for gene complexity.

Across all tissues, we observed variability by cell state, in the per cell QC metrics (fraction of mitochondrial and ribosomal reads mapped, and gene complexity per cell). To illustrate the impact of standard QC thresholds, we applied QC thresholds of 10% for the maximum mitochondrial read fraction and 500 genes detected for minimum gene complexity. A fixed cutoff of 10% mitochondrial read fraction leads to loss of parenchymal cell subsets in human kidney and testis (**Fig 2A,B**), and mouse cerebellum, and colon (**Fig S2A,B**). More broadly, mitochondrial-read-rich clusters ranged from muscle cells to tissue-parenchymal cells such as enterocytes (gut), proximal tubular cells (kidney), or sertoli cells (testis), all cell types known to have high metabolic activity and energy needs such as active transport in the kidney proximal tubule, and oxidative phosphorylation in cardiomyocytes of the heart. Even a conservative fixed cutoff of 200 genes led to loss of diverse cell subsets including immune cells such as neutrophils (**Fig S2C,D**) and neurons (**Fig 2D**). Cell cluster specific trends in percent ribosomal protein genes were also evident (**Fig 2E,F, Fig S2E,F**). Thus we demonstrate that data agnostic thresholds remove biologically relevant cells, and that QC based on these metrics must not only adapt to different tissues or samples but also to cell states and cell types.

### *ddqc*: A cell state adaptive quality control framework

To account for biological variability among QC metrics and also adapt to differences due to experimental conditions (study design, technology etc), we propose data-driven QC (ddqc, **Fig 3A**), an unsupervised, data driven, and adaptive thresholding framework for optimal capture of biological diversity. Heavily inspired by and adapting existing unsupervised approaches in scRNA-seq analysis [40], *ddqc* identifies neighborhoods of cells by graph-based clustering and performs QC on these clusters using an adaptive thresholding approach. The basic idea is that data must be partitioned by biology, and that QC must be performed on these independent partitions. Briefly, cells that pass empty droplet filters (having more than 100 genes detected and fewer that 80% of reads mapping to mitochondrial genes) are subjected to dimensionality reduction by principal component analysis, followed by nearest-neighbor graph construction and clustering to identify cell clusters with similar transcriptional states (details in **Methods**). Our approach does not rely on annotation and is driven by the density of the data. Within each such cluster, we identify “outliers” based on one- or two-sided thresholds on the QC metric of interest, defined as those cells that lie beyond a chosen number of median absolute deviations (MAD) from the cluster QC metric distribution median. Cells that pass these thresholds then enter downstream analysis.

**Figure 3:**
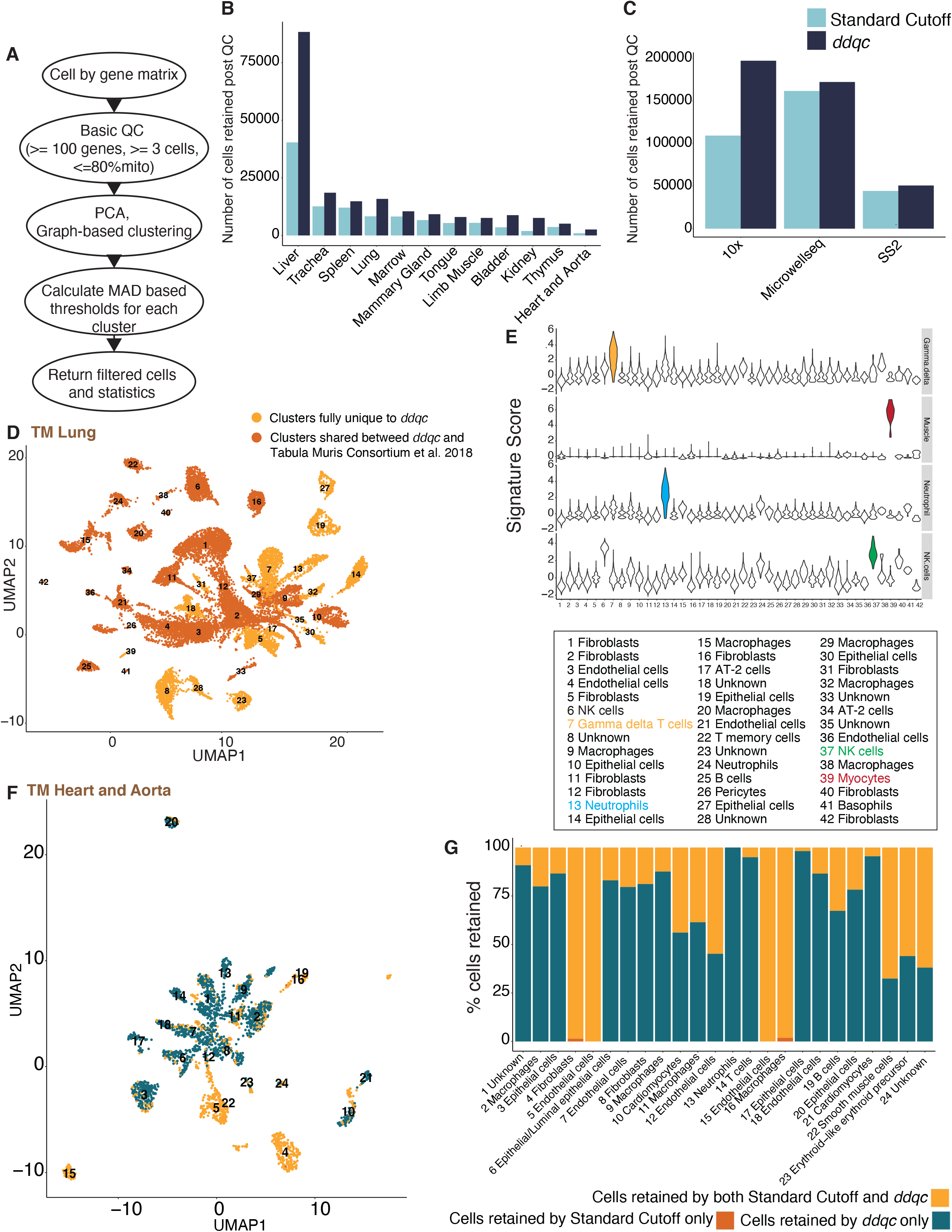
*ddqc* retains biologically meaningful cells that conventional QC filters out (A) Overview of the *ddqc* approach. (B-C) *ddqc* retains more cells when compared to the standard cutoff approach across (B) tissues in the *Tabula muris* dataset and (C) scRNA-seq data generating technologies. (D) UMAP visualization of *Tabula muris* lung cells. Colors represent whether the cells are included in the paper or uniquely retained by *ddqc*. (E) Violin plot visualization of cell type specific signature scores in average log(TPX+1). From top to bottom: muscle, neutrophil, NK cells and Gamma-delta T cells (F) UMAP visualization of joint clustering of cells retained by both *ddqc* and the standard cutoff in the mouse heart and aorta tissues. (G) Proportion of cells retained by *ddqc*, standard cutoff or both in the mouse heart and aorta tissues.

The specific downstream analysis depends on the study and biological questions of interest. For example, the next step may be integration with other data modalities (e.g. spatial data), batch effect correction or classification. If the next step is indeed conventional analysis involving clustering-based cell type identification, followed by differential gene expression, analysts may choose to start with the clustering labels that *ddqc* generates during QC (and returns as an output) to merge, re-cluster or subcluster based on their research question. *ddqc* is available as a package on Github and can be readily plugged into standard scRNA-seq analysis pipelines such as Pegasus [41] or Seurat [12]. Flexible options and exploratory plots are provided to the user for more control.

We evaluated the performance of *ddqc* on all test datasets (**Table S2**) applying adaptive QC on three QC metrics: fraction of UMIs mapped to mitochondrial genes, gene complexity and number of UMIs. For comparisons, we ran conventional QC (“standard cutoff”) on our test datasets using a fixed threshold of 10% as the maximum fraction of mitochondrial reads, and 200 as the minimum gene complexity. We then evaluated the cells that passed QC by either approach in a number of ways: ability to (1) improve power, (2) expand existing cellular taxonomies, (3) recover biologically meaningful states, and (4) discover broadly useful insights of transcriptional activity.

### *ddqc* improves power for downstream analysis when compared with conventional QC methods

We computed the number of cells retained by either *ddqc* or conventional qc and determined the breakdown by QC attributes. *ddqc* preserved more cells in comparison to conventional QC varying across datasets and biological conditions (**Table S4**). Overall, *ddqc* retained up to a median of 95.4% of input cells versus 69.4% cells using the standard cutoff approach. The higher number of cells retained by *ddqc* held across tissues (**Fig 3B**) and technologies (**Fig 3C**). Stratified by QC attributes, on average 83.19% of cells lost due to *ddqc* are due to percent mito thresholds while 6.2% are lost due to gene complexity (**Table S4**) thresholds. Thus, the higher number of cells preserved by *ddqc* provides more statistical power for downstream analysis.

### *ddqc* retains biological cell state information lost using default cutoff or data-driven approaches that do not consider biology

As *ddqc* applies QC per cluster, it helps retain several cell states of biological relevance. We illustrate the biological relevance of *ddqc* in two ways. First, using the *Tabula muris* lung dataset as a case study, we compare changes in lung cell taxonomies using *ddqc* and author-defined cutoffs. In the paper, the authors used fixed cutoffs of 500 genes for minimum gene complexity and 1000 UMIs for the minimum number of UMIs. After QC by *ddqc*, we overlaid barcode annotations (**Fig 3D**) provided by the authors [42] to define clusters with cells retained both in the paper and *ddqc*, and those exclusively retained by *ddqc* (i.e. all cells in the cluster were filtered out in the paper but retained by *ddqc*). Examining clusters exclusively retained by *ddqc*, we find various cell types of interest such as muscle cells, neutrophils, Natural Killer (NK) cells and T cells, which we validate using their known canonical signatures (**Fig 3E, Table S5**). These cell states when filtered out are not analyzed downstream. When these data are lost, we also lose the biology or insights we might learn from analyzing them. Thus, using *ddqc*, we are able to expand tissue cellular taxonomies by retaining tissue-native cell types missed by arbitrary cutoff based QC.

Next, to demonstrate that *ddqc* recovers biologically meaningful states, we proceeded to annotate the cells that passed QC using our heuristic annotation strategy. Since our annotation strategy labels cell clusters and not individual cells, we jointly clustered the cells retained by both *ddqc* and the standard cutoff QC, and then applied our heuristic clustering strategy to assign biologically relevant labels. To evaluate differences in the QC-ed cells by both approaches, we defined “uniquely retained” clusters as those that had at least 30 cell members, and 85% of cluster membership consisted of cells uniquely retained by either QC method.

No cluster was unique to the standard cutoff approach by the above definitions whereas several biologically meaningful clusters were uniquely retained by *ddqc* (**Table S6**). We describe three examples: Tabula muris Heart and Aorta **(Fig 3F,G, S4A,B),** human Olfactory Epithelial cells **(Fig S4C-F)**, the human lung **(Fig S4G,H)**. Compared to the standard cutoff method, *ddqc* retained cell subsets with low gene complexity including olfactory epithelial cells, dendritic cells, erythroid precursor cells and platelets which were filtered out by the conventional QC approach. Cardiomyocytes **(Fig S3A)** and lung muscle **(Fig 3G)** cells were mito-rich and retained in *ddqc*. The majority of cells with high mitochondrial content are diverse epithelial cells in both mouse and human. We provide a table of cell states lost when conventional methods are used across all our surveyed datasets (**Table S6**).

Finally, to compare with a data-driven approach, we ran miQC using standard settings (**Methods**) on the human olfactory epithelium and the mouse heart datasets. For the human Olfactory Epithelium, both *ddqc* and miQC retain all clusters (miQC retaining upto 95% of cells as ddqc) with *ddqc* retaining more of mito-rich olfactory epithelial cells. However, in the Tabula muris mouse heart example **(Fig S3A)**, miQC retained only 90.5% of cells as *ddqc*, completely removing the cardiomyocyte cluster. The cardiomyocyte cluster had a median of 15.178% reads mapping to mitochondrial genes, and 2427.67 as the median gene complexity, which *ddqc* retains. Cardiomyocytes are essential parenchymal cells of the heart. In both examples, miQC retained fewer cells exclusively (that *ddqc* did not), however they did not map to a missing biologically relevant cell type. Thus *ddqc* retains biologically relevant cell types that miQC filters out.

### Which cells have the least and most number of transcripts?

We next turned to other insights focusing on patterns of cell-type specific gene usage that a more biology driven QC approach preserves. Following *ddqc*, we examined trends in QC metrics (**Table S7**), to answer questions such as “which cell subsets transcribe the fewest number of genes?”. We defined cell states with low gene complexity as those with both low median number of genes detected (< 200) as well as low median %mitochondrial reads (<10%). Across 20 human studies and 163 clusters, 44 of the clusters (27%) were diverse immune cells including dendritic cells, plasma cells, T cells, Natural Killer and mast cells. Other subsets included endothelial subsets, platelets, and RBCs. Specific parenchymal cells with low gene complexity were specialized cells such as gastric chief cells (*PGA5^+^*, *PGC^+^*, *CHIA^+^*, *PGA3^+^*, *LIPF^+^*) of the stomach, cardiomyocytes (*NPPA+*, *NACA^+^*, *NACA2^+^*, *MYL2^+^*), neuronal subsets (schwann, astrocytes, neurons) of the substantia nigra, and olfactory epithelial cells. Across 4 large mouse studies and 466 clusters, 131 (28%) were immune cell clusters including 28 neutrophil (*Elane^+^*, *Prtn3^+^*, *Mpo^+^*) subsets, 27 B cells and 46 macrophage/Kupffer subsets. Endothelial (46) and erythroid (23) lineages followed. Parenchymal cells included lactating and involuting mammary gland cells, pancreatic acinar cells and diverse epithelial cells.

Next we looked at cell states with high gene complexity (> 2000 median genes, < 10% fraction mitochondrial reads). Among 318 such clusters in humans, neurons (39), and fibroblast (29) emerged as the higher ranked ones, along with epithelial cells (111). In mice, across 424 clusters, macrophages (58), fibroblasts (61), and diverse epithelial cells (104) were among the most populous subsets with high gene complexity.

### Immune cells have a high fraction of ribosomal protein content

Examining trends of ribosomal protein transcription, we defined high or low median ribosomal protein gene complexity as that with greater than 20% reads or lower than 10% reads mapping to ribosomal protein genes, and lower than 10% reads mapping to mitochondrial genes. Among 438 human clusters with high ribosomal protein gene complexity, 212 (48.4%) were immune cell subsets including 85 T cells, and 50 dendritic cell subsets. Immune cell function often requires rapid protein translation [23,43]. Other preponderant subsets were epithelial (110) and fibroblasts (43). Among 450 such clusters in mice, 241(53.6%) were annotated as immune including diverse subsets (B cell (78), macrophages (44), T cells (75)) suggesting that certain immune states may have high translational activity and need for ribosomal protein genes.

Neurons (20.6%) were a large fraction of human cell states with lower ribosomal protein gene complexity. In mouse, cell states with low ribosomal protein gene complexity included diverse epithelial and immune cells, fibroblasts and endothelial cells. Thus a more context-focused QC approach such as *ddqc* can enable us to recapitulate and study fundamental patterns in cell-type specific gene expression and associated function.

## Discussion

Cell quality-control remains an essential step in scRNA-seq data analysis, however conventional approaches apply arbitrary filters on defined QC metrics without accounting for the biological context. The standard practice among published papers is largely a data-agnostic arbitrary threshold-based QC. We have demonstrated (**Fig 1-2**) that not accounting for the underlying biological heterogeneity at the level of cell states during QC can lead to loss of relevant biological insights (including important cell types) as well as reduced statistical power for downstream analysis. However, identifying cell types and cell states is a time-consuming process requiring either well-annotated training sets or involves the manual and subjective task of cell-state annotation. The field of single-cell biology is still in the early stages of building experimentally validated and reproducible ontologies of cell states. To overcome these challenges, we present an unsupervised approach *ddqc* that leverages clustering to identify transcriptionally similar cellular neighborhoods (approximating broad cell types) and performs adaptive QC on these clusters.

We observe limitations of our approach: (1) *ddqc* applies adaptive thresholds on each cluster, and hence, we are likely to lose some good quality cells due to inherent spread of the cluster data distribution. (2) While in most cases, *ddqc* retains clusters that are biologically meaningful, in some cases, *ddqc* may retain cells (**Table S4**) with high percentages of mitochondrial genes that may be a mix of biology and technical artifacts. These clusters when sub-clustered do not always represent bimodal distributions (**Fig S3**), rather a gradation and there is no perfect way to assess the right cutoff. Such cells are usually subsets of larger neighborhoods of biologically meaningful cells that reflect true metabolic stress due to the biological condition studied. In the current version of *ddqc*, removal of such cells has been left to the analyst after examination via Exploratory Data Analysis (EDA) in the context of the biology of the study, and during downstream analysis. We believe QC should be iterative and to help empower the user, ddqc provides detailed statistics for all cells that pass or fail adaptive QC.

*ddqc* provides several advantages relative to conventional cutoff or biology-agnostic data driven approaches. First, it retains more cells than standard or data-driven QC approaches leading to more power for downstream analysis. Second, the additional cells retained by *ddqc* are biologically meaningful thus increasing the potential for further biological discovery. Such biological insights include retaining a diversity of cell types with extreme value QCs and rare cells, as well as uncovering study-specific metabolic and physiological programs that may dictate changes in these common QC metrics. Further investigation of retained cell states may provide insights into the underlying biological processes. Finally, we examine cells lost by conventional QC to add insights into questions of fundamental interest in biology such as parsimony in total gene usage or transcription. Our analysis has revealed interesting biological observations in terms of overall transcriptional diversity of cell states, as well as ribosomal protein gene expression. In summation, we propose a biology-centered and iterative approach to cell quality control that retains cell states of critical biologically relevance often removed by conventional QC.

## Code Availability

*ddqc* is available as a GitHub package along with a tutorial: https://github.com/ayshwaryas/ddqc For R users, a compatible package is available here: https://github.com/ayshwaryas/ddqc_R

## Data availability

All datasets used in this manuscript are publicly available. Supplementary tables 1 and 2 provide the information for accessing the datasets.

## Acknowledgements

We are immensely grateful to Aviv Regev for her mentorship, helpful suggestions and resources. We gratefully acknowledge Oana Ursu, Sean Simmons and Matan Hofree for critical review of the manuscript. We acknowledge authors of several published studies who kindly responded to data requests or questions. We thank Angela Pisco and James Webber (Tabula Muris), Kyle Joseph Travaglini (Human Lung dataset), Jonathan Manning (EBI) and Dmitry Velmeshev (ASD snRNA-seq) for sharing datasets. We thank the MIT PRIMES program, the Broad Institute Academic Affairs, Carrie E Wager, Elana Gonsalves, Anna Greka, Vijay Kuchroo, Ana Anderson and members of the Regev lab.

Earlier iterations of this work were presented at the Women in Statistics and Data Science Conference in 2018, and the MIT PRIMES Fall Conference in 2019.

## Author contributions

AS conceived the project, wrote the pilot version of *ddqc*, performed analysis, interpreted results and wrote the paper. MA co-developed *ddqc* with input from AS, performed analysis and contributed to the Methods section of the manuscript. YY and BL helped with integration of *ddqc* into Pegasus. All authors read the manuscript, provided input and have approved it for submission.

---

**Supp Fig 1:**
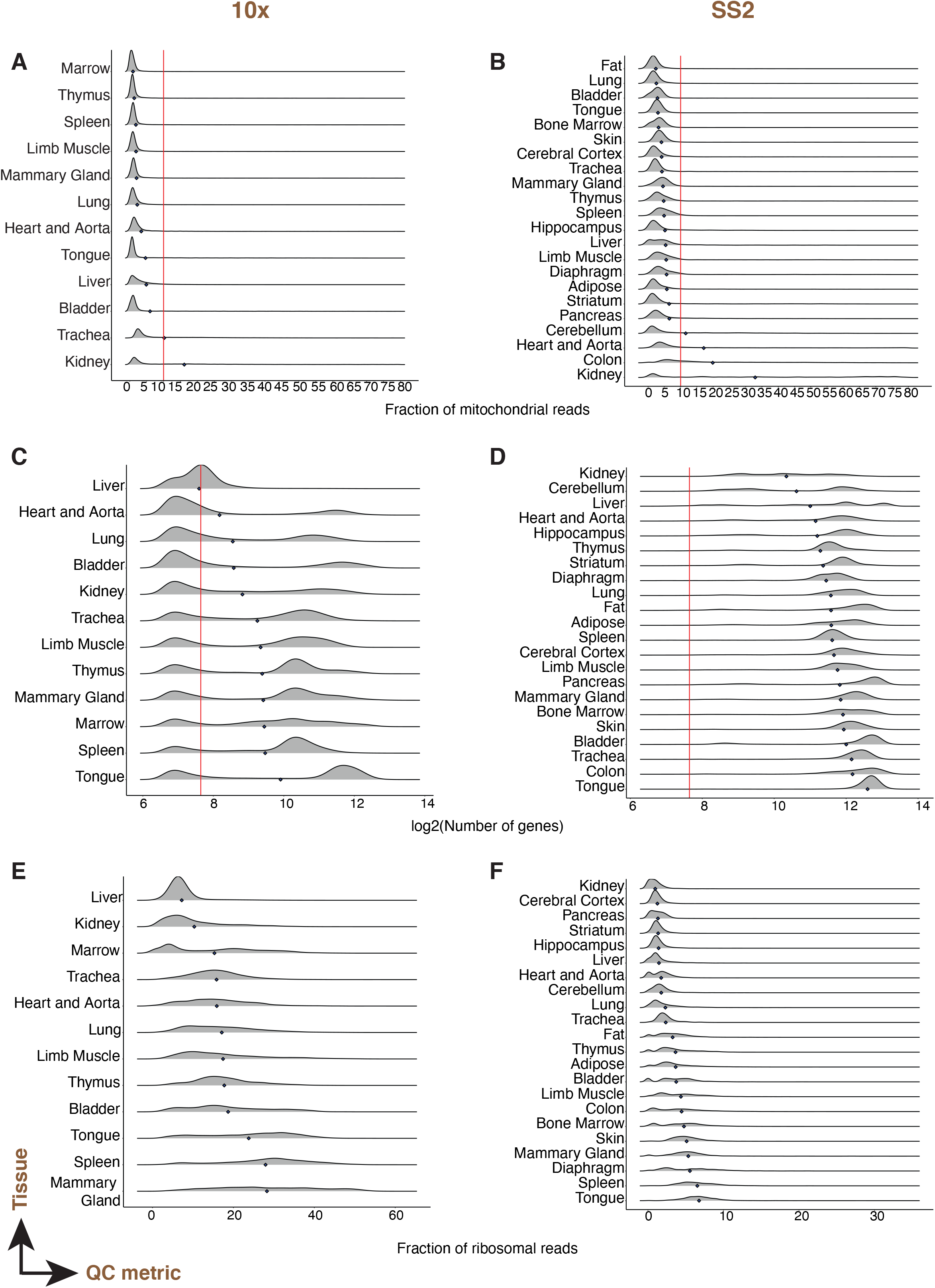
QC metrics vary by tissue: Fraction of mitochondrial reads (A,B), gene complexity (C,D) and percentage of ribosomal protein genes (E,F) per cell across mouse tissues and technologies. Various mouse tissue scRNAseq datasets generated in the *Tabula muris* project by 10X droplet-based (A,C,E) and Smart-seq2 (SS2; B,D,F) plate-based technologies. Each row in a panel is a density curve with the mean represented by a blue diamond. Red lines indicate conventional threshold values set at 10% for percentage of mitochondrial reads, and 200 for gene complexity.

**Supp Fig 2:**
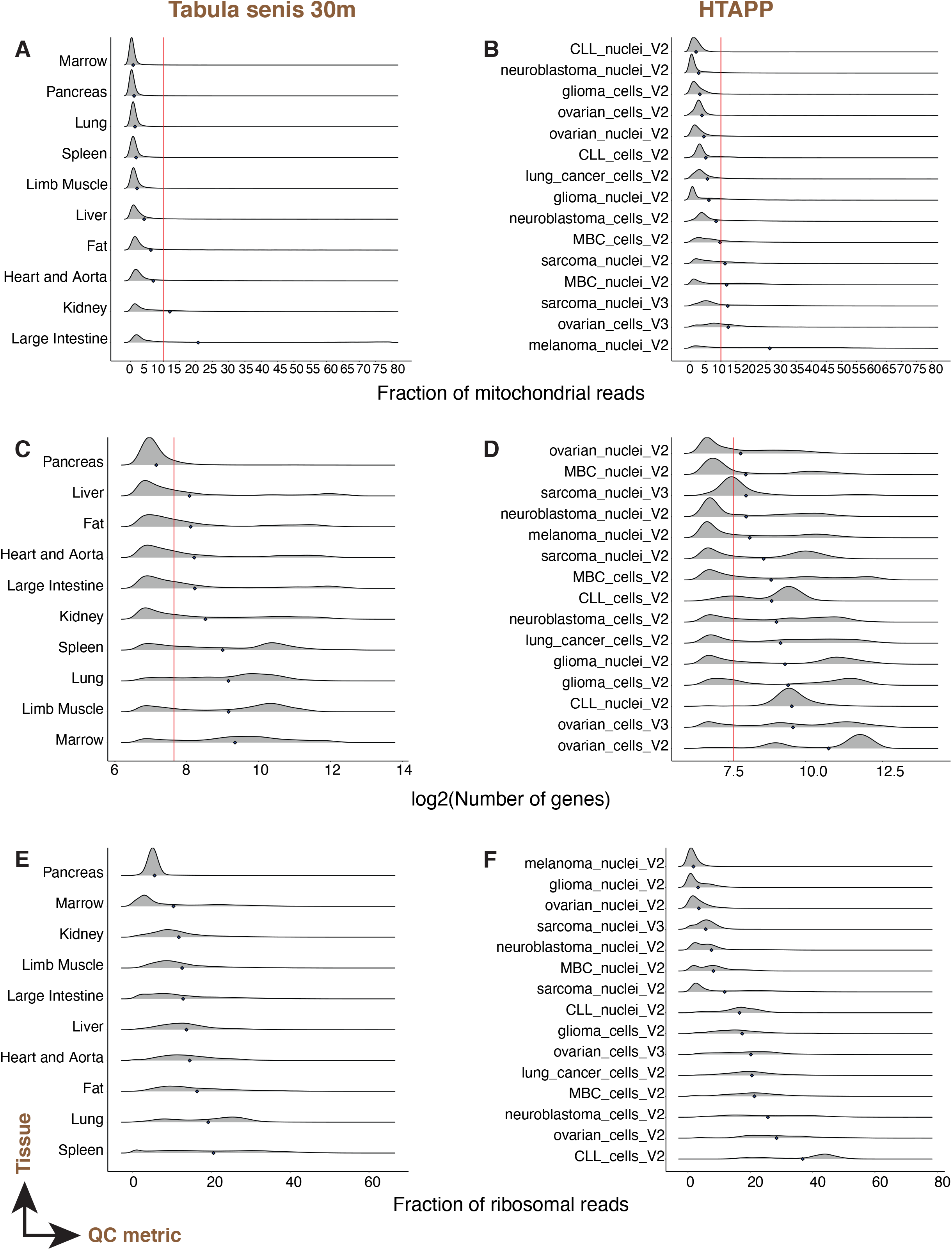
QC metrics vary by tissue: Fraction of mitochondrial reads (A,B), gene complexity (C,D) and percentage of ribosomal protein genes (E,F) per cell across mouse tissues and cancers. Various mouse tissue scRNAseq datasets generated in the *Tabula senis* project (30 months) (A,C,E) and the human tumor atlas pilot project (HTAPP; B,D,F) by 10X droplet-based. Each row in a panel is a density curve with the mean represented by a blue diamond. Red lines indicate conventional threshold values set at 10% for percentage of mitochondrial reads, and 200 for gene complexity.

**Supp Fig 3:**
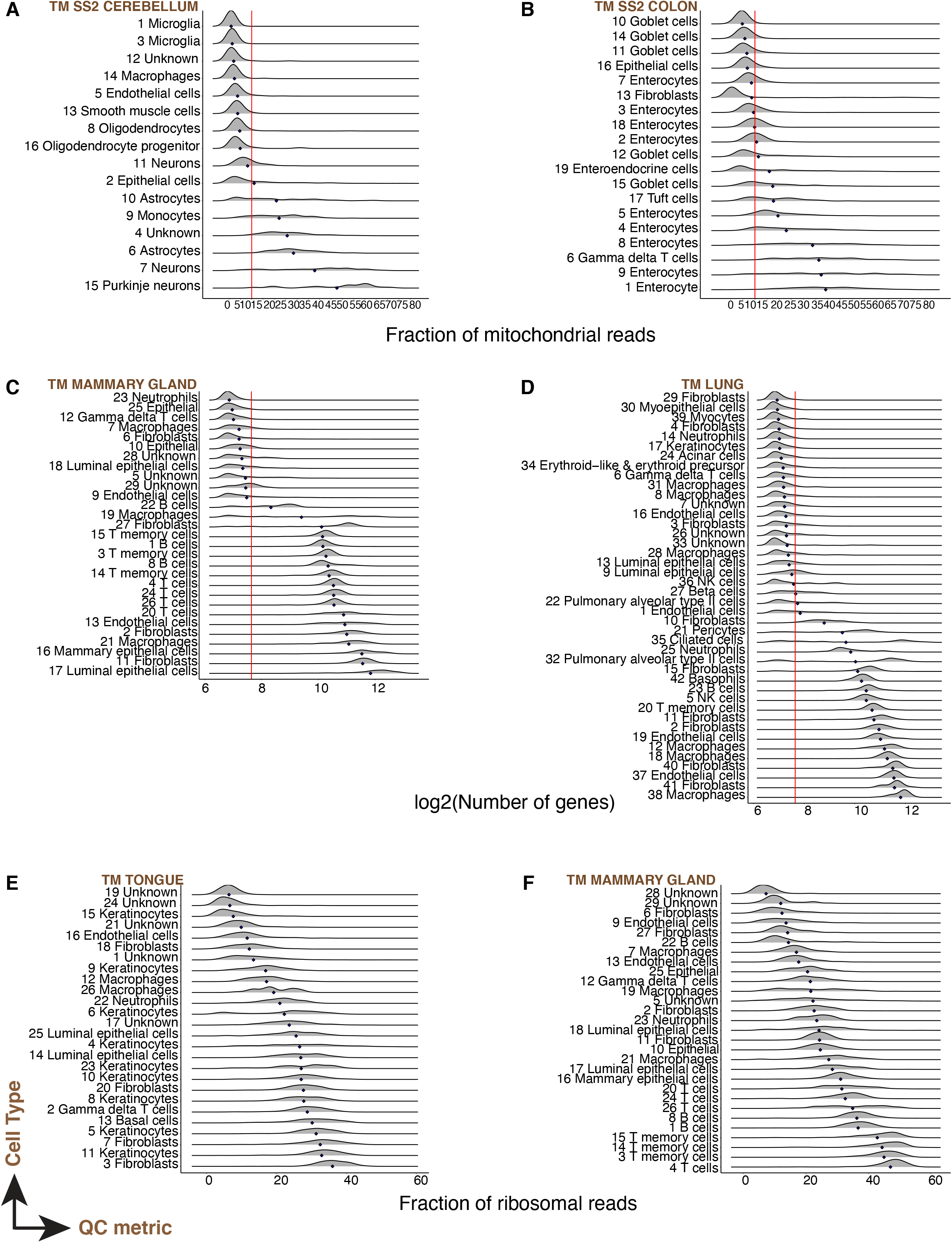
QC metrics vary by cell-type: Fraction of mitochondrial reads (A,B), gene complexity (C,D) and percentage of ribosomal protein genes (E,F) per cell across cell types of various mouse tissues: Cerebellum (A), colon (B), mammary gland (C), lung (D), tongue (E) and Lung (F). All scRNA-seq data was generated using the 10X droplet-based technology. Each row in a panel is a density curve with the mean represented by a blue diamond. Red lines indicate conventional threshold values set at 10% for percentage of mitochondrial reads, and 200 for gene complexity.

**Supp Fig 4:**
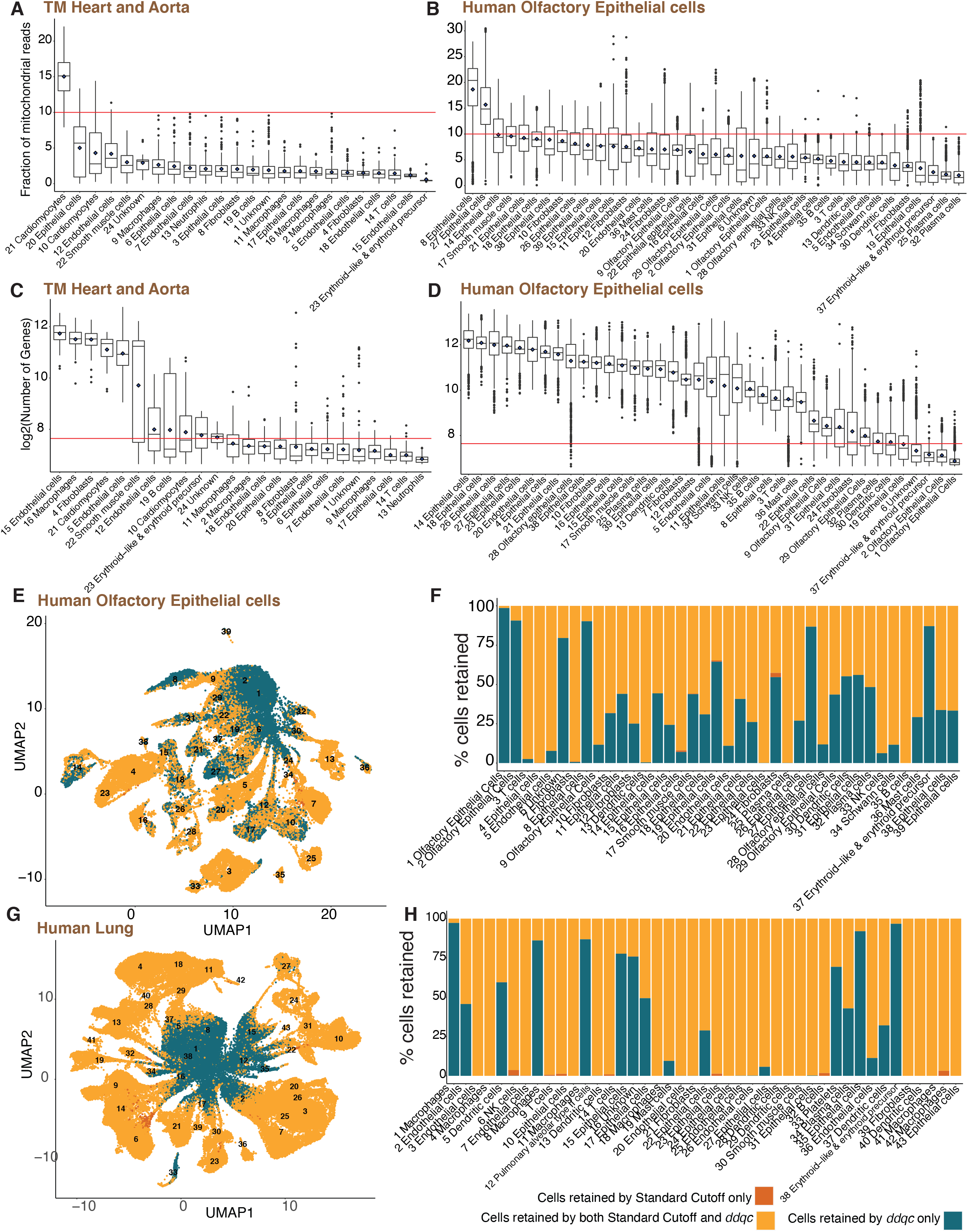
*ddqc* retains cell states of biological relevance. Boxplot visualization of the fraction of reads mapping to the mitochondria, and the gene complexity across cell types in the (A,C) mouse heart and aorta and (B,D) human olfactory epithelium. (E,G) UMAP visualization of joint clustering of cells retained by both *ddqc* and the standard cutoff in (E) human olfactory epithelium and (G) human lung. Proportion of cells retained by *ddqc*, standard cutoff or both in (F) human olfactory epithelium and (H) human lung.

**Tables**

Supp Table 1: Table of papers included in the QC survey

Supp Table 2: Table of datasets with details of source tissue, samples, publications.

Supp Table 3: Details of annotations used in the evaluation of the automatic annotation strategy

Supp Table 4: QC Statistics returned by *ddqc* and standard cutoff

Supp Table 5: Tabula muris lung: comparisons with published annotations and signatures

Supp Table 6: Table of clusters unique to either *ddqc* or standard cutoff

Supp Table 7: Trends in cell types with extreme QC metrics

## Methods

### QC survey

We conducted a survey of 107 single-cell and single-nucleus RNA sequencing papers published between 2017-2020. Papers included in the survey were collated either from Twitter posts, searches on Google or from the scRNAseq database [44]. For each paper, we recorded Quality Control (QC) strategy from the “Methods” section into Table S1. Additional information was also recorded for each paper:

- Year published
- Organism
- Tissue of Origin
- sequencing technology
- Analysis Software
- Preprocessing software

QC was classified into the following categories:

- QC to remove low-quality cells and genes by QC metric

- number of counts
- number of genes
- percent of mitochondrial transcripts
- percent of ribosomal transcripts
- number of cells in which gene is present
- QC to remove empty droplets
- QC to remove doublets/multiplets
- QC to account for Ambient/Background RNA

We categorized the papers based on which type of QC each paper used for a particular metric. These categories were:

- Data-agnostic fixed threshold - QC removed all cells with a metric above/below a certain number (for example keep all cells with <10% mitochondrial transcripts)
- Multiple fixed thresholds - several fixed thresholds thresholds for different samples
- Data-Driven study-level threshold - QC threshold was determined from the data (for example, keep all cells with a number of genes within 2 SDs from the median)
- Custom - QC that was very specific for the particular paper
- No filtering - no filtering based on this metric was done

Summary of the QC survey and QC methods are documented in Table 1 and the Results section.

### Datasets

We downloaded publicly available mouse (n=5) and 32 human (n=32) (Table S2) single-cell (scRNA-seq) or single-nucleus (snRNA-seq) RNA sequencing datasets. We restricted our study to droplet- (10X Genomics), MicrowellSeq and plate-based (SmartSeq2) technologies from various tissues.

We downloaded data at the level of gene counts after preprocessing (genomic reference alignment and gene-level quantification) but prior to any quality control (QC). However, many datasets in public repositories were already filtered using cutoffs, or were aligned to reference genomes with missing genes. In some cases, we were able to contact study authors (Tabula Muris) and get the unfiltered expression matrices. Links to the unfiltered datasets used can be found in Table S2. Our dataset search was agnostic to the computational preprocessing methodology or genome reference version used.

### Basic preprocessing

For all analyses, we start with loading the unfiltered or raw cell-by-gene matrix stored either in the mtx, csv, txt, or h5ad format. Basic QC or initial filtering is conducted to remove poor quality cells: cells with less than 100 genes or with more than 80% of mitochondrial transcripts and genes present in less than 3 cells are removed. The basic QC step is also important for computational efficiency as otherwise, we may have on the order of a million or more barcodes incase of droplet based scRNA-seq.

### ddqc

We propose an adaptive thresholding method to perform quality control at the level cell types, thus taking into account differences between them. The first step of this method is to cluster the cells using standard scRNAseq analysis preprocessing and clustering steps. We assume that in each cluster cells are of the same or closely related cell types with shared biological properties. In each cluster, we expect outliers - cells with the number of UMI counts, number of genes, or percent of mitochondrial transcripts significantly different from the cluster average. We assume that those differ in quality from other cells in their cluster, and remove them by calculating a cutoff for each cluster based on median absolute deviation and a predefined parameter. We chose the median absolute deviation (MAD) to be a more robust statistic to define outlier thresholds instead of the zscore which assumes normality, or IQR which is less permissive.If the cell has a value higher (percent.mito) or lower (n_counts, n_genes) than 2 MADs from the median in its cluster, this cell will be filtered out; all remaining cells will be sent for downstream analysis. If the cluster *ddqc* threshold was bigger than 200 n_genes, or lower than 10% mito, we would set it to 200 or 10 respectively.

*ddqc* uses preprocessing and clustering functions provided by the Pegasus (https://pegasus.readthedocs.io/) for the Python package: https://github.com/ayshwaryas/ddqc. An R package using functions in Seurat is also available: https://github.com/ayshwaryas/ddqc_R.

Our pipeline starts with a loading of the unfiltered cell-by-gene matrix stored either in mtx, csv, txt, or h5ad format.

- Basic QC was conducted to remove obvious bad quality cells: cells with less than 100 genes or with more than 80% of mitochondrial transcripts using the functions *qc_metrics* and *filter_data* (*subset* in R).
- Normalization is performed using the function *NormalizeData* (*NormalizeData* in Seurat): normalize the feature expression measurements for each cell by the total expression, multiply by a scale factor (10,000), and log-transform the result to get log(TPX+1) values.
- We find the top 2000 highly variable genes using the function call *highly_variable_features* (*FindVariableFeatures* in Seurat). We scale the expression matrix of highly variable genes: shift the expression of each gene so that the mean expression across cells is 0 and scale the expression of each gene so that the variance across cells is 1, (In pegasus scaling is part of *pca*, in Seurat *ScaleData*)
- Next, dimensionality reduction is performed using principal component analysis (PCA) using *pca* (*RunPCA*) with the number of principal components set at 50.
- Graph-based clustering of cells was performed by first building the k-nearest neighbor graph setting K=20 [45], and then the Louvain algorithm for clustering [46] or community detection with the resolution set at 1.4 using the functions *neighbors* (*FindNeighbors*) and *louvain* (*FindClusters*) functions.
- Then we iterate through each of QC metrics to determine the cutoff values:

- First we create a true/false numpy array (vector in R) that would represent whether the cells have passed ddqc
- For each cluster we find lower (for n_counts and n_genes, otherwise set to negative infinity) and upper (percent mito, otherwise set to positive infinity) cutoff (median ± 2 * MAD)

◾ For number of genes: If lower cutoff is less than 200 genes, it would be set to 200 (by default)
◾ For percent mito: if upper cutoff is more than 10 percent, it would be set to 10 (by default)
- Finally, f the cell is outside the bounds defined by cutoffs, it would be marked as false in the ddqc array
- We do an *AND* operation between all ddqc metric-specific arrays. Cells that are marked as true in this array have passed ddqc and are retained for downstream analysis

In the Pegasus and Seurat workflows, in addition to returning the filtered object, ddqc returns a pandas dataframe with the following information for each cell:

- True/False value that indicates whether the cell passed the ddqc
- Cluster number that was assigned to this cell in the initial clustering
- For each QC metric:

- The metric itself
- Lower cutoff (cluster median - 2 cluster MAD) for this metric for the cell’s cluster. If there is no cutoff, this field will be equal to None
- Upper cutoff (cluster median + 2 cluster MAD) for this metric for the cell’s cluster. If there is no cutoff, this field will be equal to None
- True/False value that indicates whether the cell passed the ddqc for the given metric

In addition, the ddqc workflow displays two box plots: one shows percent mito by cluster with red line at 10 percent that indicates the standard fixed threshold for percent mito, and the other shows log2 of n_genes by cluster with red line at 200 genes (7.64 in log2-scale) that indicates the most common fixed threshold for number of genes.

### Automatic cell-type annotation

We automated the task of mapping cell types to clusters using the PanglaoDB cell type gene expression signatures as the reference dataset. Using the PanglaoDB cell type: marker mappings, cell type labels were assigned for each cluster as follows:

1. We computed cluster specific differentially expressed genes (DGE) by testing for genes differentially expressed in the cluster of interest vs all else. For the testing, we used the default differential expression test used in Seurat for the R version or Pegasus for the Python version.
2. We filtered the DGE to retain those genes with at least a log fold change of > 0.25, percent expressed in the cluster of interest > 25%, and p value < 0.05
3. We iterated through each cluster to assign cell type scores as follows:

a. First, we iterated through the filtered DGE of the current cluster to check for matches in PanglaoDB.

i. If there was an entry that the gene indicates for a particular cell type, the average log fold change of that gene was added to the score of the cell type.
ii. Only cell types annotations which included at least three such marker genes were retained
b. The cluster was assigned the cell type annotation with the highest score. Otherwise, the cell type would be stated as Unknown.

We note that the accuracy of our method is contingent on the accuracy of markers in the PanglaoDB dataset which would get updated on a regular basis. The PanglaoDB markers database doesn’t have enough genes for certain cell types, which causes them to be wrongly identified (For example Macrophages are often labeled as Dendritic cells). For examples in Figures 2 & 3, annotations were manually verified.

### Automatic cell-type annotation accuracy assessment

In order to assess the accuracy of our cell type annotations method we have compared the results of automatic annotations with the annotation provided by the publisher of the dataset, if such annotation was provided. Datasets where the authors provided annotations included the Human Tissue Atlas; human adipose (inhouse annotated), heart, and lung; *Tabula muris* (10x), *Tabula muris* (Smartseq-2), *Tabula senis* 10x 24 and 30 month. The accuracy was calculated using the steps below.

1. First, we annotate the clusters after only empty droplet filtering. We do it by mapping the annotation that is the most frequent among the cells of the cluster. If most of the cells don’t have an annotation, the cluster will be marked as unknown.
2. For accuracy analysis, we are only keeping the clusters that had an annotation (not unknown) and where at least 75% of cluster cells had that annotation.
3. For the comparison, we have established a number of pairs of the annotations that we are considering to be the same **(Table S3).** Some of those pairs are just different in naming (example NK cells VS Natural Killer cells), and others were validated by marker genes to be more accurately defined using our strategy.
4. Then, we count the number of clusters with a mismatch between automatic annotation and annotation provided by the publisher. If the annotation pair is included in the table from step 3, it will not be counted as a mismatch. After that, we compute the accuracy percentage.

The tables of the same cell types, mismatches, exact numbers and breakdown by the dataset is provided in **Table S3**.

### Comparison of ddqc with author-provided annotations

We have compared *ddqc* with author-provided quality control in tabula muris tissue:

1. First, the author-provided annotations were downloaded from figshare (https://figshare.com/articles/dataset/Single-cell_RNA-seq_data_from_microfluidic_emulsion_v2_/5968960?file=13088039).
2. Then we calculated the percent of cells exclusive to *ddqc* in each cluster after ddqc filtering **(Table S6).** It was calculated by taking the number of cells whose barcodes were not present in author annotations (which means they were not included by the author for final analysis) and dividing it by the total number of cells in the cluster.
3. To verify the automatic annotation for clusters with high percent (100%) exclusive, we have computed signature scores for each of the clusters (using the “pegasus.calc_signature_score” function) with cell type markers **(Figure 3e).** You can find the signature genes in the **(Table S6).**
4. We have also generated UMAP plots with cells colored based on percent exclusive of their cluster. We had 2 categories: fully exclusive to ddqc or shared with paper. **(Figure 3D)**

### Comparison of *ddqc* to the standard cutoff method

We compared *ddqc* with the standard cutoff or static threshold method (default in most pipelines) as a control, and only basic QC for reference:

1. *ddqc* using the same steps as described in the *ddqc* section for loading the data and filtering.
2. Standard Cutoff or Static threshold (cells with number of genes less than 200 and mitochondrial transcripts percent higher than 10% are removed regardless of filtering)
3. No additional filtering (done for reference)

First, we evaluated the retained cells in all the three approaches independently by graph based clustering, followed by differential gene expression using *de_analysis* function and UMAP visualization using *umap* the function. Also, additional statistics were recorded for future analysis (Information about clusters and cells). Exploratory data analysis (EDA) was performed by generating summary plots including boxplots, joyplots, and colored UMAP plots.

Next, for comparisons, we performed joint clustering as follows:

1. After QC was performed, each barcode is assigned a label which indicates if it was filtered or retained by each method. Possible options are: retained by all methods, retained by ddqc only, retained by Cutoff only, Neither (removed by both cutoff and ddqc)
2. Barcodes that were marked as “neither” were removed
3. All remaining barcodes were clustered (as above), and visualized using UMAP.
4. Both cluster and filter labels were used to color the UMAPs for exploratory data analysis. Barplots were also generated per cluster to visualize the distribution of each cluster by cell retained in each method.
5. DGE was performed on the clusters to assign cell identity, and to identify cell-types lost by single-threshold QC.

These plots helped to demonstrate differences between static threshold and *ddqc* by highlighting clusters of cells that were kept by one method, but lost by another.

#### Unique Clusters

To demonstrate differences between static threshold and *ddqc,* we determined how many meaningful “unique” clusters *ddqc* retained. A “unique” cluster was defined as a cluster with at least 30 cells, and with at least 85% of its cells retained only by *ddqc*, but filtered out by cutoff method. The presence of unique clusters indicates that a population of very similar cells was almost entirely filtered by one method, thus suggesting that potentially some cell types were exclusive only to the other method. This helped to demonstrate the advantage of ddqc over a static threshold since it had many more unique clusters than the static threshold method had. More detailed examples are provided in the results section.

### Comparisons with miQC

At the time of testing, miQC was installed in R from GitHub using the command “remotes::install_github(“greenelab/miQC”, build_vignettes = TRUE)”. miQC was run on the test datasets using the standard steps as described in the vignette: https://github.com/greenelab/miQC/blob/main/vignettes/miQC.Rmd. Comparison was performed by examining the intersection of miQC retained barcodes with those retained by *ddqc*, leveraging the annotations in the *ddqc* results.

### Trends table

We determined trends in QC metrics by iterating through all ddqc clusters in all tissues and recording the clusters which satisfy one of the following criteria to a corresponding table:

- Median number of genes lower than 200
- Median number of genes higher than 2000
- Median percent mito higher than 10
- Median percent ribo lower than 10
- Median percent ribo higher than 20

### Visualization and plotting

Boxplots, joyplots and violin plots for each QC metric was generated in R using the *ggplot2* and *ggridges* packages. For the tissue summary plots (Figure 1) only basic QC was performed, and then the QC metrics plotted stratified by tissue. For cell-type summary plots (Figure 2), graph-based clustering was performed after basic QC. A horizontal red line for boxplots and violin plots, and vertical line for joyplots were added to illustrate standard cutoff thresholds (10% for mitochondrial transcripts percent, 200 for number of genes).

All analysis tasks were performed on the Broad Institute High-Performance Computing Cluster.

